# JAK/STAT mediated insulin resistance in muscles is essential for effective immune response

**DOI:** 10.1101/2023.10.04.560867

**Authors:** Ellen McMullen, Lukas Strych, Lenka Chodáková, Amber Krebs, Tomas Dolezal

## Abstract

The metabolically demanding nature of immune response requires nutrients to be preferentially directed towards the immune system at the expense of peripheral tissues. We study the mechanisms by which this metabolic reprograming occurs using the parasitoid infection of *Drosophila* larvae. To overcome such an immune challenge hemocytes differentiate into lamellocytes, which encapsulate and melanize the parasitoid egg. Hemocytes acquire the energy for this process by expressing JAK/STAT ligands upd2 and upd3, which activates JAK/STAT signaling in muscles and redirects carbohydrates away from muscles in favor of immune cells. Absence of upd/JAK/STAT signaling leads to an impaired immune response and increased mortality. We demonstrate how JAK/STAT signaling in muscles leads to suppression of insulin signaling through activation of ImpL2, the inhibitor of Drosophila insulin like peptides. We reveal the crucial function of muscles during immune response and show the benefits of insulin resistance as an adaptive mechanism that is necessary for survival.

## Introduction

Immune response is a highly energy demanding process, requiring an efficient and rapid supply of nutrients to immune cells. Upon activation, immune cells increase their metabolic demand, more than doubling their glucose consumption (preprint: Kazek *et al*., 2023). This requires a systemic metabolic shift to ensure the immune system has sufficient energy to overcome the infection (Bajgar *et al*., 2015). According to the concept of selfish immunity, resources can be redirected to the immune system by inducing insulin resistance in non-immune tissues (Straub, 2014). Insulin resistance is predominantly studied in a pathological context (Gurzov *et al*., 2016; Lourido *et al*., 2021) and its adaptive function during the acute immune response is largely unexplored (preprint: Krejčová *et al*., 2021).

We examine the mechanisms that facilitates the redistribution of nutrients during infection of *Drosophila melanogaster* by parasitoid wasp *Leptopilina boulardi* and how this metabolic shift is essential for effective immune response. *Leptopilina boulardi* lay their eggs inside *Drosophila* in the early third instar larval stage (Russo *et al*., 1996). Upon recognition of the wasp egg, hemocytes, *Drosophila* immune cells, differentiate into lamellocytes, which encapsulate and melanize the wasp egg (Nappi and Christensen, 2005). If successful immune response is mounted, the *Drosophila* will pupate and reach adulthood without major setbacks; however, if it is insufficient the parasitoid larva will continue to grow and kill the *Drosophila* larvae.

The primary objective of *Drosophila* larvae is to feed and grow; meaning metabolites are supplied to developing tissues, such as muscles as well as the fat body, the *Drosophila* equivalent of the liver and adipose tissue. However, upon infection, as metabolic demand increases in the immune system, carbohydrates are directed away from peripheral tissues in favor of immune cells (Bajgar *et al*., 2015; Bajgar, Krejčová and Doležal, 2021). This requires the regulation of metabolic signaling on an organismal level and cross talk between different organs and tissues to coordinate nutrient supply. Previous studies have shown that expressions of cytokines unpaired 2 (upd2) and unpaired 3 (upd3) are upregulated within a few hours in hemocytes of infected larvae (Yang *et al*., 2015). Through binding to the domeless receptor, upd ligands activate JAK/STAT signaling in muscles, which is required for the differentiation of lamellocytes and efficient immune response to parasitoid wasps. JAK/STAT signaling is a highly conserved regulator of immune response (Yang and Hultmark, 2017; Yu *et al*., 2022), and has been shown to have important immunological and metabolic roles throughout the body, including the muscle, fat body and gut (Yu *et al*., 2022).

JAK/STAT signaling is known to play an important role in metabolic regulation and growth (Shin *et al*., 2020); it has also been shown to suppress insulin signaling in contexts such as host wasting in cancer models (Ding *et al*., 2021). Therefore, we hypothesized that the release of upd cytokines from hemocytes during wasp infection can suppress insulin signaling in muscles, thereby reducing their nutrient consumption, leaving more nutrients available for immune response. Both JAK/STAT and insulin signaling are required in muscles during larval development for proper feeding and consequently effective immune response (Yang and Hultmark, 2017). However, the interaction between JAK/STAT and insulin signaling during infection has not been firmly established. Here we show that upd2 and upd3 activate JAK/STAT signaling and suppress insulin signaling in muscles, as demonstrated by a reduction in muscle glycogen stores. Absence of upd2 or upd3 significantly impairs immune response; likewise, muscle specific knockdown of *STAT* results in insufficient lamellocyte production and decreased survival.

JAK/STAT signaling directly induces the expression of Ecdysone-inducible gene L2 (*ImpL2*), an insulin binding protein (Terry *et al*., 2006; Amoyel *et al*., 2016). We illustrate that, as is the case with adult *Drosophila, ImpL2* plays an important role in the cross talk between immune cells and the periphery, in terms of nutrient distribution during immune response (preprint: Krejčová *et al*., 2021). In the case of this manuscript, we focus on the communication that takes place between muscle tissue and the immune system. Silencing of *ImpL2* in muscles negatively impacts immune response, while over expression of *ImpL2* in muscles offers a partial rescue in lamellocyte production and larval survival in *upd* mutants.

Previous studies have showed the role of the upd/JAK/STAT axis in certain pathologies such as host wasting in cancer (Ding *et al*., 2021). Here, we show the importance of this mechanism in immune response. We shed light on the cross talk that occurs between the immune system and muscles during immune response. We demonstrate the suppression of insulin signaling in muscle tissue during infestation allows for the redirection of carbohydrates away from muscles through the ‘selfish signaling’ of immune cells. This prioritization of the immune system is fundamental for *Drosophila* survival.

## Results

### Carbohydra televels change upon infection

As seen in our previous work (Bajgar *et al*., 2015), in the wild type situation, upon infection, circulating trehalose levels drop (Figure 1 A) as trehalose is broken down into glucose molecules. Glycogen stores in muscles are also reduced (Figure 1C), as carbohydrates are directed away from muscles in favor of the immune system. This leads to an increase in glucose levels as the animals reach a state of hyperglycemia offering a readily available energy source for hemocytes (Figure 1B). Prioritization of the immune system in this manner leads to a developmental delay of the larvae, as well as a decreased movement (Figure 1D), as muscles have less energy to function. However, in *upd* mutants, this upd mediated metabolic shift is absent, therefore energy sources are not reallocated from muscles to the same extent, resulting in no significant reduction in either circulating trehalose (Figure 1A) or muscle glycogen levels (Figure 1C). This is further demonstrated by the fact that infected *upd* mutant animals grow and move at the same rate as their uninfected counterparts (Figure 1D). Interestingly *upd* mutant animals have lower circulating glucose levels compared to control animals, but glucose levels still increase upon infection (Figure 1B). Decrease in muscle glycogen stores and an increase in glucose in the circulation in wildtypic animals demonstrates the metabolic shift that occurs during infection. The absence of such change in *unpaired* mutants emphasizes the importance of this signal to mediate this redirection of energy during immune response.

**Figure 1:**
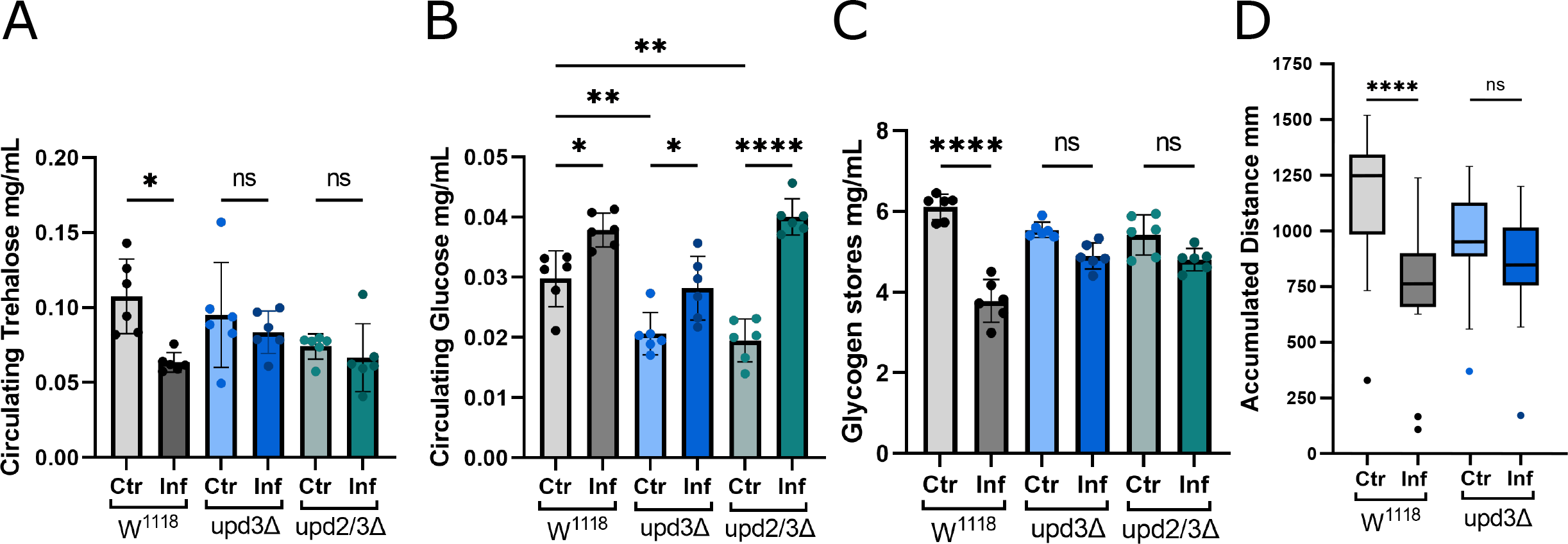
In upd mutants carbohydrates are not redirected away from muscles upon infection A: Comparison of circulating trehalose levels of control animals and upd mutants in non-infected and infected conditions (N=2, n=6). B: Circulating Glucose levels of upd mutants compared to controls in non-infected and infected larvae (N=2, n=6). C: Level of glycogen stores in muscle tissue in non-infected and infected animals (N=2, n=6). D: Larval locomotion: Accumulated distance traveled by control animals and upd3 mutants, with and without infection (N=2-3, n=18-24). A-C: Bars represent mean values, dots represent biological replicates. D: boxes represent mean values, dots represent individual larvae. ns: no significant difference, * p ≤ 0.05, ** p ≤ 0.01, **** ≤ 0.0001. N represents individual experiments, n represents biological replicates.

### upd2 and upd3 play a key role in immune response

Unpaired cytokines are responsible for the initiation of JAK/STAT signaling by binding to the domeless receptor (Morin-Poulard, Vincent and Crozatier, 2013). Loss of upd2 or upd3 leads to impaired lamellocyte production and therefore an inadequate immune response to wasp infection (Yang *et al*., 2015) (Figure 2A). This results in a reduced survival rate of the *Drosophila* larvae (Figure 2B), as there are insufficient lamellocytes to encapsulate and melanize the wasp egg. As such, the wasp larvae outcompetes the *Drosophila* and continues to develop and grow, eventually killing the *Drosophila* larvae and emerging from the *Drosophila* pupal case. We show that upd2 and upd3 are necessary for sufficient lamellocyte production; without which immune response is severely diminished, as is the survival rate of the *Drosophila* larvae.

**Figure 2:**
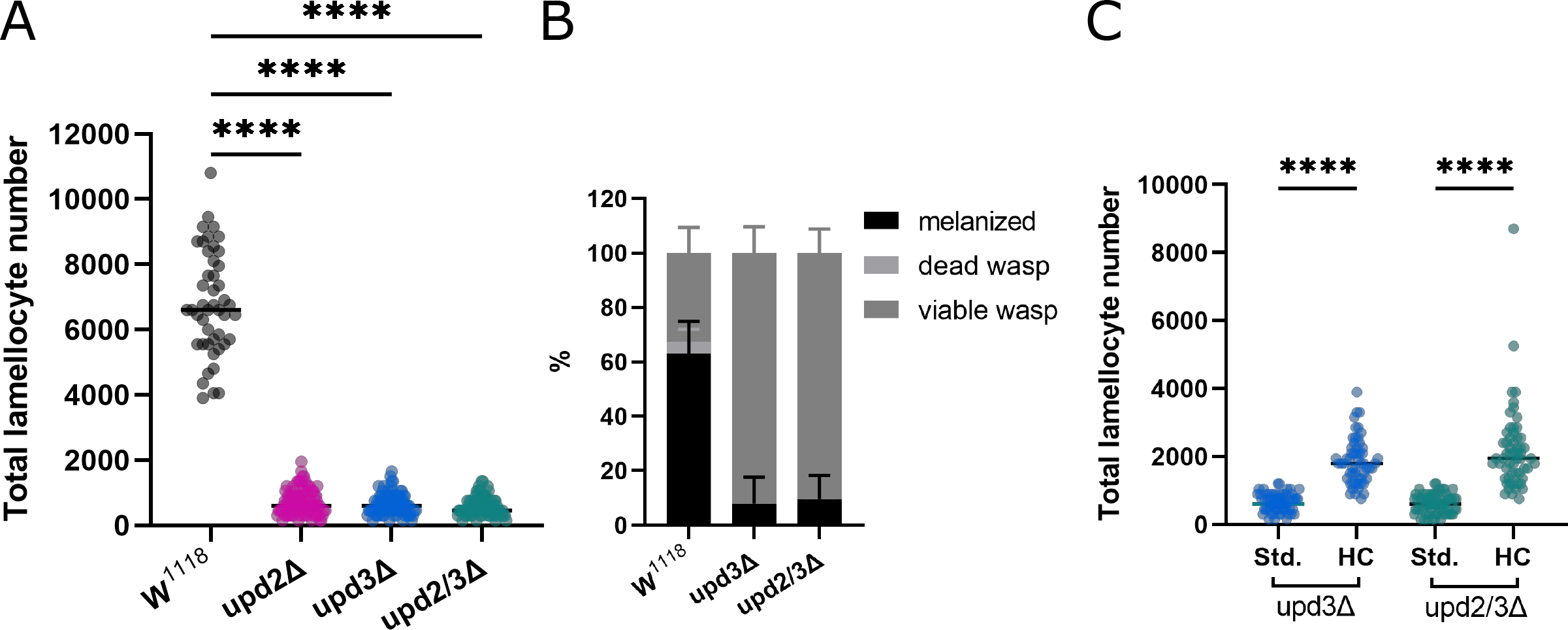
The role of upd in immune response A: Lamellocyte number in upd mutants compared to control animals (N=3, n=46-99), each dot represents number of lamellocytes in an individual larvae. B: Resistance to wasp infection (N=3, n=43-45), there is a significant difference in number of melanized wasps between upd3Δ and control and upd2/3Δ and control animals (p = ≤ 0.0001). C: Lamellocyte number in upd mutants on high carbohydrate food (N=3, n=56-86), each dot represents number of lamellocytes in an individual larvae. **** ≤ 0.0001. N represents individual experiments, n represents biological replicates.

### Glucose supplementation ameliorates impaired immune response

The introduction of a high carbohydrate diet partially rescues lamellocyte differentiation and resistance against wasp infection seen in *updΔ* animals. When *updΔ* larvae are transferred to a diet supplemented with 10% glucose at the point of infection, they are able to produce more lamellocytes than those on a standard diet (Figure 2C). This data shows that adequate access to nutrients is essential for effective immune response.

### JAK/STAT signaling in muscles is necessary to mediate immune response

As loss of upd results in an inadequate immune response, we investigated whether disrupting JAK/STAT signaling by other means would have a similar affect. By utilizing the Gal4 UAS Gal80 system (thermosensitive Gal80 inhibiting Gal4), we performed muscle specific knockdown of STAT in the muscles of late second instar larvae, just prior to infection. This allowed us to circumvent issues that occur by knocking down STAT throughout development (Yang and Hultmark, 2017). Knock down of STAT in muscle tissues, by RNAi, phenocopies the reduction in lamellocyte number seen in *upd* null mutants (Figure 3A). With fewer lamellocytes STAT knock down animals befall a similar fate to *updΔ* animals, as survival post infection is greatly reduced compared to the *eGFP* controls (Figure 3B). This gives further evidence of the importance of upd/JAK/STAT signaling in immune response and shows that this signal is essential specifically in muscles.

**Figure 3:**
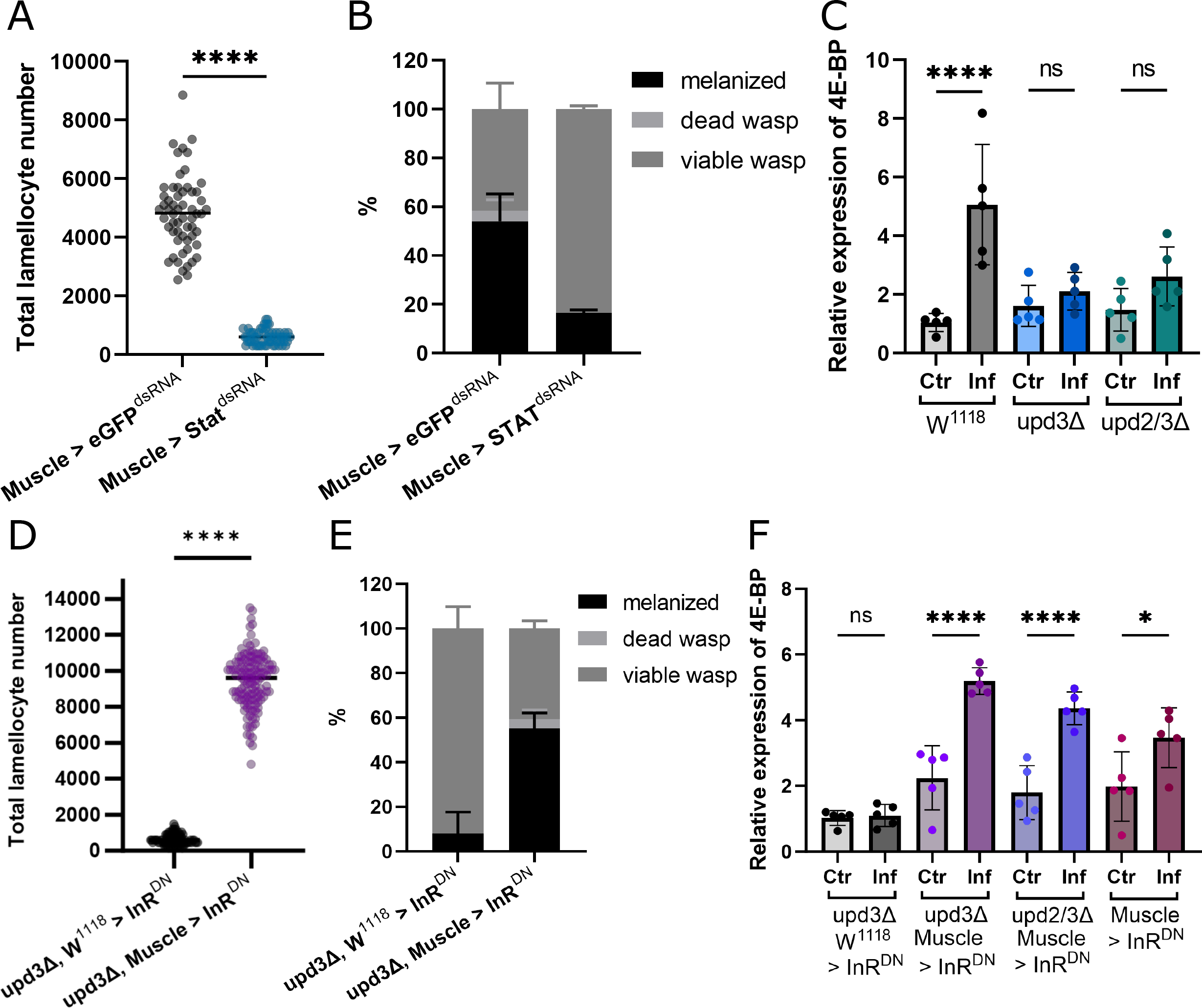
JAK/STAT signaling in muscles is mediated by insulin signaling A: Number of lamellocytes in control and Stat^dsRNA^ animals (N=3, n=58-59), dots represent the number of lamellocytes of individual larvae. B: Comparison of resistance to wasp infection between control and Stat^dsRNA^ animals (N=3, n=43-55), the statistical difference in number of melanized wasps between the two genotypes p = ≤ 0.0001. C: Relative expression of 4E-BP in non-infected and infected control and upd mutant animals (n=5), bars represent mean values with each dot showing a biological replicate. D: Lamellocyte number of upd3 mutant animals and animals of the same genetic background expressing InR^DN^ specifically in muscles (N=3, n=97-133), dots represent the number of lamellocytes of individual larvae. E: Comparison of wasp resistance between upd3 animals with and without the expression of InR^DN^ in muscles (N=3, n=46-49), statistical difference in number of melanized wasps p = ≤ 0.0001. F: Relative expression of 4E-BP in non-infected and infected control animals and upd mutants expressing InR^DN^ in muscles (n=5), bars represent mean values with each dot showing a biological replicate. ns: no significant difference, * p ≤ 0.05, **** ≤ 0.0001. N represents individual experiments, n represents biological replicates.

### Hematopoiesis is not affected by upd/JAK/STAT manipulation

Despite the striking difference in lamellocyte production observed in the multiple genetic manipulations of the upd/JAK/STAT signaling pathway we see no such change in plasmatocytes number (Figure S1A-F). This strongly suggests that disruption of upd/JAK/STAT signaling limits only the differentiation into lamellocytes rather than hematopoiesis itself, as there is insufficient energy allocated to the immune system during wasp infection.

### Insulin signaling in muscles is essential for effective immune response

To determine whether the diminished immune response observed in *upd3Δ* and *upd2/3Δ* animals was, in fact, due to insulin signaling we looked at the expression of FOXO target 4E-BP in the muscle tissues of these larvae. FOXO is a known regulator of insulin signaling (Jünger *et al*., 2003); therefore, we used 4E-BP levels as a read out to see whether insulin signaling in muscles is altered upon infection. As expected, the relative expression of 4E-BP significantly increases in control animals, indicating an increase in FOXO signaling upon infection (Figure 3C). This is likely due to suppression of insulin signaling in muscles, as a means to divert metabolites away from muscles in favor of the immune system as metabolic demand increases during infection. However, we observed no such increase in 4E-BP in *upd* null mutants (Figure 3C), as the signaling to mediate this metabolite diversion is silenced. These results suggest that insulin signaling plays a role in metabolic regulation during infection.

### Expression of InR ^DN^ can overcome loss JAK/STAT signaling in terms of immune response

We showed that expression of 4E-BP is increased upon infection, which indicates a link to insulin signaling. We propose that upd/JAK/STAT signaling regulates insulin signaling in muscles, which in turn is vital for effective immune response. Blocking this signal by knockout of upd or knockdown of JAK/STAT results in a limited immune response, therefore we wanted to see if we could rescue immune response by suppressing insulin signaling through other means. To do so we expressed InR^DN^, the dominant negative version of insulin receptor, in muscle tissues of *upd* null mutants, again using Gal4 UAS Gal80 system to induce expression just prior to infection. We show that suppression of insulin signaling in muscles by the expression of InR^DN^ can rescue the reduction in lamellocyte number seen in *upd* null mutants (Figure 3D) and consequently survival rates of these animals (Figure 3E).

When insulin receptor dominant negative is expressed in the skeletal muscle of *upd* mutants, we see higher levels of the metabolic regulator 4E-BP even without infection (Figure 3F). This is likely due to the artificial suppression of insulin signaling in these animals and it demonstrates that expression of InR^DN^ indeed leads to suppression of insulin signaling in muscles. Upon infection 4E-BP expression increases significantly as metabolic demand increases and more energy is required to mount an immune response. Together, these results suggest suppression of insulin signaling in peripheral tissues, such as muscles, is necessary to facilitate metabolic reprograming during infection.

### Suppression of insulin signaling is not due to lower Dilp expressions

*Drosophila* insulin-like peptides (Dilps) play a role in many functions of development, growth and ageing, especially in terms of metabolic regulation (Slaidina *et al*., 2009; Wang, Karpac and Jasper, 2014). Therefore, we looked how levels of specific Dilps change due to infection. We measured the expression of Dilp2, Dilp3 and Dilp5 in the brains of larvae 28 hpi (Figure S1 A) and Dilp6 in the fat body of these same larvae (Figure S2 B). Upon infection the expression of Dilp3 and Dilp5 increase significantly, with Dilp2 also following that trend (Figure S1 A). Likewise, expression of Dilp6 in the fat body tissue increases after infection (Figure S1 B). Based on this we suggested that the metabolic changes that we observe during infection are due to insulin resistance in peripheral tissues, such as muscles, rather than an insulin deficiency.

### Expression of ImpL2 in muscles is necessary for immune function

JAK/STAT signaling has been shown to directly trigger the expression of ImpL2 (Terry *et al*., 2006; Amoyel *et al*., 2016). The Ecdysone-inducible gene was shown to regulate systemic metabolism by reducing systemic insulin signaling (Honegger *et al*., 2008). To explore the relationship of ImpL2 in our system we knocked down ImpL2 in a muscle specific manner using RNAi. As predicted, loss of ImpL2 in muscles lead to a reduction in lamellocyte production compared to control animals (Muscle > eGFP^dsRNA^) (Figure 4A). As a result, the survival rate of these *Drosophila* larvae was also impacted, with fewer *Drosophila* successfully overcoming the wasp invasion (Figure 4B).

**Figure 4:**
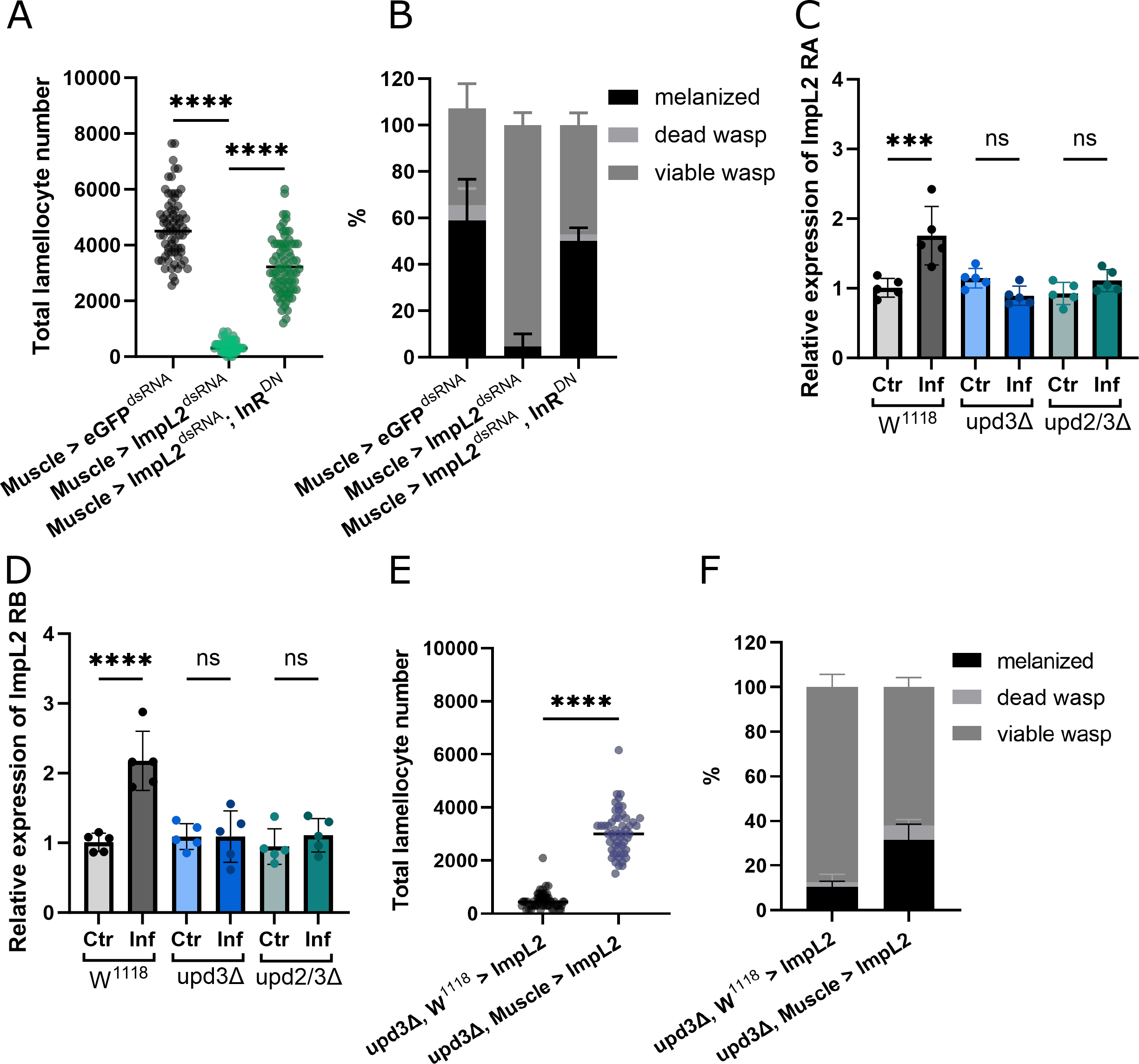
ImpL2 mediates suppression of insulin signaling A: Lamellocyte number in animals with muscle specific knockdown or ImpL2^dsRNA^ compared to eGFP^dsRNA^ controls and and Muscle > ImpL2dsRNA InRDN animals 22 hpi (N=3, n=45-80), each dot represents a single larvae. B: Percentage of viable drosophila and wasp larvae in Muscle > eGFP^dsRNA^, Muscle > ImpL2^dsRNA^ and Muscle > ImpL2dsRNA InRDN animals after wasp infestation (N=3, n=51-55), there is no significant difference between number of melanized wasps in Muscle > eGFP^dsRNA^ and Muscle > ImpL2^dsRNA^, InR^DN^; the number of melanized wasps in Muscle > ImpL2^dsRNA^ is significantly fewer (p = ≤ 0.0001) than the other genotypes. C: Expression of ImpL2 RA in control and upd animals with and without infection (n=5), bars represent the mean values, each dot represents a biological replicate. D: Expression levels of ImpL2 RB with and without infection in control and upd animals (n=5), bars represent the mean values, each dot represents a biological replicate. E: Number of Lamellocytes in upd3 animals with (upd3Δ, Muscle > ImpL2) and without (upd3Δ, W^1118^ > ImpL2) the overexpression of ImpL2 in muscle tissue (N=3, n=53-58), each dot represents a single larvae. F: Resistance to wasp infection of upd3Δ, W^1118^ > ImpL2 and Muscle > upd3Δ, Muscle > ImpL2 animals (N=3, n=53), p = 0.0283 for the difference between number of melanized wasps. ns: no significant difference, *** p ≤ 0.001, **** ≤ 0.0001. N represents individual experiments, n represents biological replicates.

**Figure 5:**
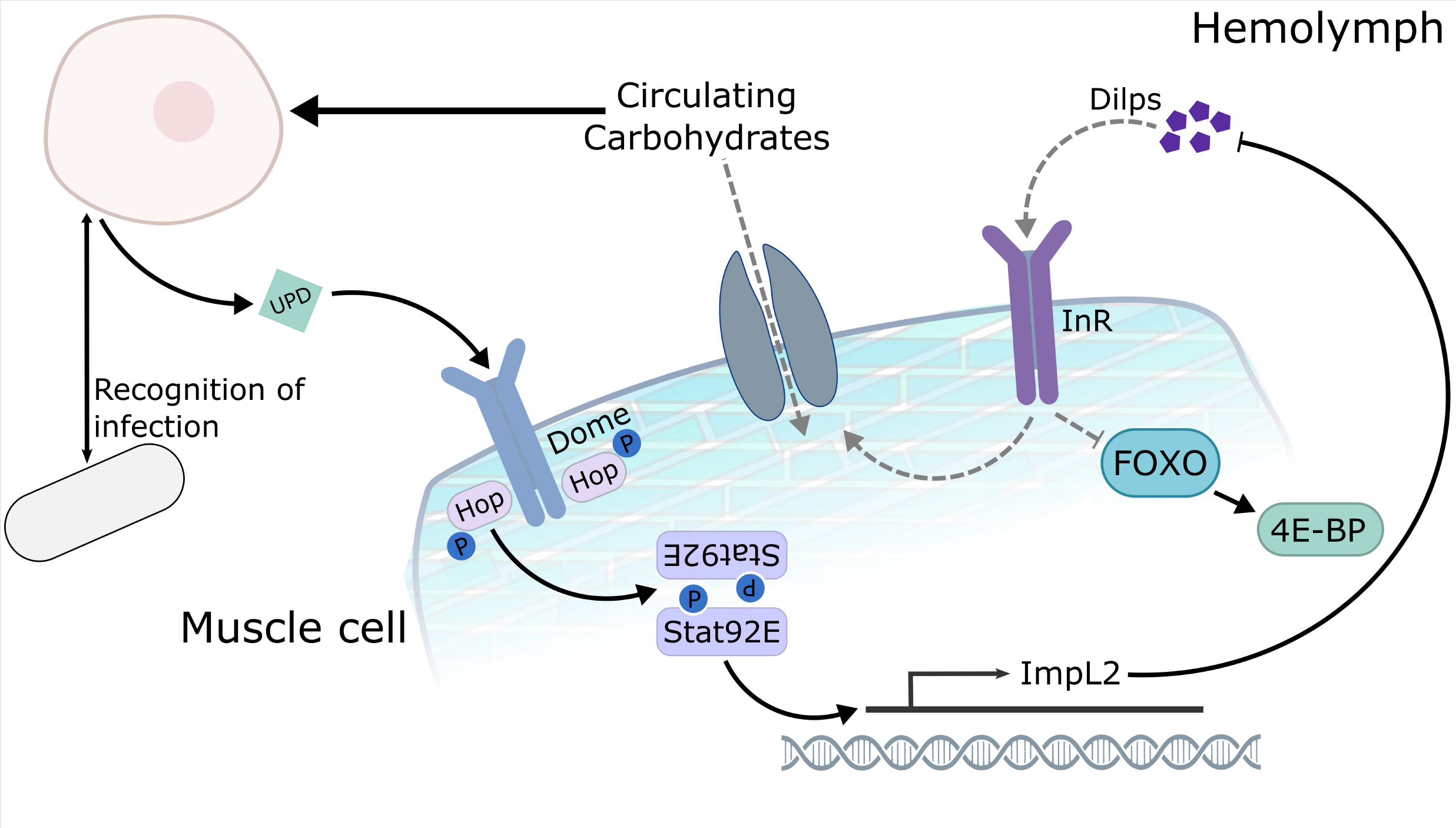
Schematic representation of signaling pathway during infection During infection, upd/JAK/STAT triggers the expression of ImpL2 in muscles leading to the local suppression of insulin signaling. This allows for the redirection of nutrients away from muscles, in favor of the immune system. Disruption in the upd/JAK/STAT signaling pathway, through genetic manipulation means insulin resistance in muscles does not occur and therefore the immune system does not acquire sufficient nutrients to mount an effective immune response.

To consider the link between upd and ImpL2 we looked at how expression levels of ImpL2 change upon infection in muscles. In the wild type situation levels of both ImpL2 RA (Figure 4C) and ImpL2 RB (Figure 4D) increase upon infection as more ImpL2 is produced in order to reduce insulin signaling and therefore free up metabolites that can be directed towards the immune system. However, no such increase is observed in *updΔ* animals (Figures 4C and 4D). Additionally, expression of ImpL2 actually decreases upon infection in hemocytes (Figure S2 C). This supports our previous findings that the ‘selfish’ immune signal, and therefore metabolic shift upon infection, is not occurring to the same extent in *updΔ* larvae. Moreover, these results highlight that release of ImpL2 from muscles plays a role in immune response.

### Expression of InR ^DN^ rescues impaired immune response in ImpL2 knockdown animals

As expression of InR^DN^ in the muscle of *upd* null mutants was able to improve immune response, and survival, we wanted to see if the same could be achieved in *ImpL2* knockdown animals. Therefore, we simultaneously suppressed ImpL2 in muscles, while also expressing InR^DN^. In this situation we saw a significant increase in lamellocyte production in Muscle > ImpL2^dsRNA^; InR^DN^ animals compared to Muscle > ImpL2dsRNA animals (Figure 4A). This is reflected in a higher survival rate, similar to the rate observed in control animals (Muscle > eGFP^dsRNA^) (Figure 4B). This shows that the suppression of insulin signaling in muscles via ImpL2 expression is indeed a key regulator of immune response.

### Over expression of ImpL2 in muscles offers improved immune response in upd mutants

Knock down of ImpL2 in muscles leads to a reduction in lamellocyte number and chance of survival. Our previous results suggest that upd executes its function via the induction of ImpL2 in muscles; which is missing in *updΔ* larvae. Therefore, we wanted to see what happens if we overexpress ImpL2 in *updΔ* animals. By expressing ImpL2 in the background of *upd3Δ* animals, we see a partial rescue in lamellocyte number as well as survival rate compared to *upd3Δ* which do not express ImpL2 (*upd3Δ, w*^*1118*^ *> ImpL2*) (Figures 4E and 4F). This shows the importance of ImpL2 signaling in muscles as a regulator of immune response.

## Discussion

The health of any animal relies on many factors, including the tight regulation of metabolic processes. Disruption in metabolic homeostasis can be detrimental, leading to reduced fitness and even death. Throughout the lifespan of an animal metabolic demand changes due to development, ageing or other challenges (Bajgar *et al*., 2015; Son *et al*., 2019). Utilizing the power of *Drosophila* and a parasitoid infection model, we unravel inter-organ communication that occurs between the immune system and muscles during immune response. Here we show that the initiation of JAK/STAT signaling, by upd, in muscles leads to the suppression of insulin signaling during infection, which is essential for survival.

Mounting an effective immune response is extremely costly in terms of energy. Therefore, suppression of systemic metabolic processes, coupled with the reallocation of metabolites during infection is a conserved process across the animal kingdom (Wang, Luan and Medzhitov, 2019). In the case of parasitoid invasion in *Drosophila* larvae, nutrients are redirected away from non-immune tissues to provide energy for the activation and differentiation of immune cells (Bajgar *et al*., 2015). Production of lamellocytes is necessary for the encapsulation and neutralization of the invading wasp egg (Tokusumi *et al*., 2009). Without sufficient lamellocytes, survival of the *Drosophila* is significantly decreased.

Here, we give further insights into the metabolic reprogramming that occurs during infection, namely the suppression of insulin signaling and redirection of energy stores away from muscles. This leads to an increase in circulating glucose levels and ultimately the utilization of these sugars by immune cells.

Additionally, in infected animals circulating trehalose, the primary sugar in *Drosophila*, is broken down into glucose, as a ready source of carbohydrates for immune cells. In combination, this provides sufficient fuel to the immune system to mount an effective immune response. However, this process of decreased carbohydrate supply to muscles leads to developmental delay (Bajgar *et al*., 2015) and reduction in movement of the larvae; but ultimately causes no lasting consequences.

Such metabolic changes require the cross talk between the muscle tissue and the immune system, mediated by cell signaling. A promising candidate for such an interaction is JAK/STAT signaling. Previous studies have alluded to a link between JAK/STAT and insulin signaling, however the exact mechanism at play during immune response has remained elusive (Yang and Hultmark, 2017). By utilizing a ubiquitous Gal80 to suppress Gal4 expression during development, we were to gain greater understanding of the relationship between JAK/STAT and insulin signaling by looking at this interaction exclusively during immune response. We show that JAK/STAT mediated insulin resistance in muscles is a key regulator of carbohydrate allocation during immune response. Blocking this signal, either through silencing of upd2 or upd3 ubiquitously, or specifically in muscles using STAT^dsRNA^, leads to a reduction in lamellocyte number and therefore survival of the larva, as the immune system does not receive sufficient energy resources. The supplementation of glucose in the diet of *upd* mutants allows for the production of more lamellocytes (partial rescue), as more carbohydrates are available and can be utilized by the immune cells. Likewise, the expression of InR^DN^ in the muscle tissue of *upd* mutants offers a rescue in both number of lamellocytes and resistance to parasitoid wasp. The addition of InR^DN^ offers an alternative signal to suppress insulin signaling, therefore restoring the metabolic switch needed to fuel the immune system.

Suppression of insulin signaling in tissues has been observed in pathologies, such as cancer (Ding *et al*., 2021; preprint: Liu *et al*., 2023). Here we demonstrate the importance of host mediated insulin resistance as an adaptive mechanism, crucial for survival. We show that JAK/STAT signaling in muscles, as initiated by upd cytokines, results in the suppression of insulin signaling within muscles. These findings show the benefits of the suppression of insulin signaling in selective tissues and emphasize that coordinated insulin resistance is in fact advantageous.

Upd signaling is important for suppression of insulin signaling in muscles. The release of upd from hemocytes (Agaisse *et al*., 2003; Yang and Hultmark, 2017) (Figure S2 C) provides further evidence of the selfish immune system. The theory of the selfish immune system was previously demonstrated by the release of adenosine from immune cells to obtain more nutrients (Bajgar *et al*., 2015). Here we show the role of upd/JAK/STAT as another key mechanism to secure a metabolic advantage for the immune system. It appears that immune cells trigger JAK/STAT mediated insulin resistance in muscles, thereby granting themselves privileged access to nutrient supply. In the control situation, relative expression of the FOXO target 4E-BP in muscles increases four fold in infected animals. However, no significant increase is observed in *upd* mutants after infection. When insulin signaling is diminished in muscles through the expression of InR^DN^ in *upd* mutants basal 4E-BP expression is considerably higher than that of the control, increasing further in infected animals. This data supports previous findings that 4E-BP serves as a metabolic break during challenging conditions (Teleman, Chen and Cohen, 2005). We suggest that upd2 and upd3 are key regulators in metabolic regulation in parasitoid infestation, without which infected animals are unable to overcome this immune challenge.

We shed more light on the role of IGFBP7 homolog ImpL2 during immune response. ImpL2 is a negative regulator of insulin signaling (Honegger *et al*., 2008). It has previously been show that the hemocytes of adult *Drosophila* release ImpL2 as an immune response (preprint: Krejčová *et al*., 2021). Additionally, tumor cells express ImpL2 to suppress insulin signaling in host tissues (Ding *et al*., 2021; preprint: Liu *et al*., 2023). However, we show here for the first time, in *Drosophila* larvae, that activation of JAK/STAT signaling in muscles leads to the local expression of ImpL2, which in turn suppresses insulin signaling.

Expression of ImpL2 in muscles themselves facilitates the mobilization of nutrients to be utilized by the immune system as required. This is demonstrated by the suppression of ImpL2 in muscles, resulting in fewer lamellocytes, and thus decreased survival rate. On the other hand, when ImpL2 is over expressed in muscles of *upd* mutants there is a partial, yet significant rescue in both production of lamellocytes and resistance to wasp infestation. The increased expression of ImpL2 upon infection in muscles of control animals is further evidence of the antagonistic role of ImpL2 in insulin signaling. As was observed in *upd* null mutants, muscle specific expression of InR^DN^ leads to a rescue in both lamellocyte number and survival rate, once again demonstrating the role of ImpL2 in insulin resistance during infection. These findings also demonstrate a novel mechanism of ImpL2 release from the muscles causing insulin resistance within the same tissue. Although we cannot exclude that ImpL2 has a systemic effect, by suppressing insulin signaling in other tissues, our results show that in terms of metabolic signaling during immune response muscles do the heavy lifting.

Systemic changes in metabolism are important in times of nutrient scarcity or increased metabolic demand (Smith *et al*., 2018). It is this ability to adapt that is crucial for survival. We demonstrate here how upd/JAK/STAT mediated metabolic reprograming allows immune cells to preferential acquire carbohydrates during immune response. We also demonstrate the role of ImpL2 as a key player in regulation of insulin signaling and mobilization of metabolites during immune response. Moreover, we show that suppression of insulin signaling in muscles by upd/JAK/STAT/ImpL2 signaling is a fundamental host response to immune challenge, without which the animal is unlikely to overcome the infection.

## Materials and Methods

### Fly Stocks

Fly stocks were maintained at room temperature, crosses were performed at 18 °C and transferred to 25 °C 24 hours prior to infection. All experiments we performed on male larvae.

Flies were raised of a diet of cornmeal (80 g/l), agar (10 g/l), yeast (40 g/l), saccharose (50 g/l) and 10% methylparaben.

For selected experiments, larvae were transferred to a high carbohydrate diet: (80 g/l), agar (10 g/l), yeast (40 g/l), glucose (180 g/l) and 10% methylparaben at point of infection.

Following fly stocks were obtained from Bloomington stock center:

*w*^*1118*^, *w*^*1118*^ upd2Δ (BL55727), *w*^*1118*^ upd3Δ (BL55728), *w*^*1118*^ upd2/3Δ (BL55729), Stat^dRNA^: P{UAS-Stat92E.RNAi}1 (BL26899), ImpL2^dsRNA^: P{TRiP.HMC03863}attP40 (BL55855), InR^DN^: P{UAS-InR.K1409A}3

(BL8253), Muscle driver: P{GawB}how[24B] (BL1767), *w*^*1118*^; P{w[+mC]=tubP-GAL80[ts]}2 (BL7017) was recombined with P{GawB}how[24B], eGFP: y[1] sc[*] v[1] sev[21]; P{y[+t7.7] v[+t1.8]=VALIUM20-EGFP.RNAi.1}attP2 (BL41556)

ImpL2^s.UAS^: *UAS-ImpL2* (*UAS-s*.*ImpL2*; FBal0249386) was gift from Dr. Hugo Stocker

### Primers stocks

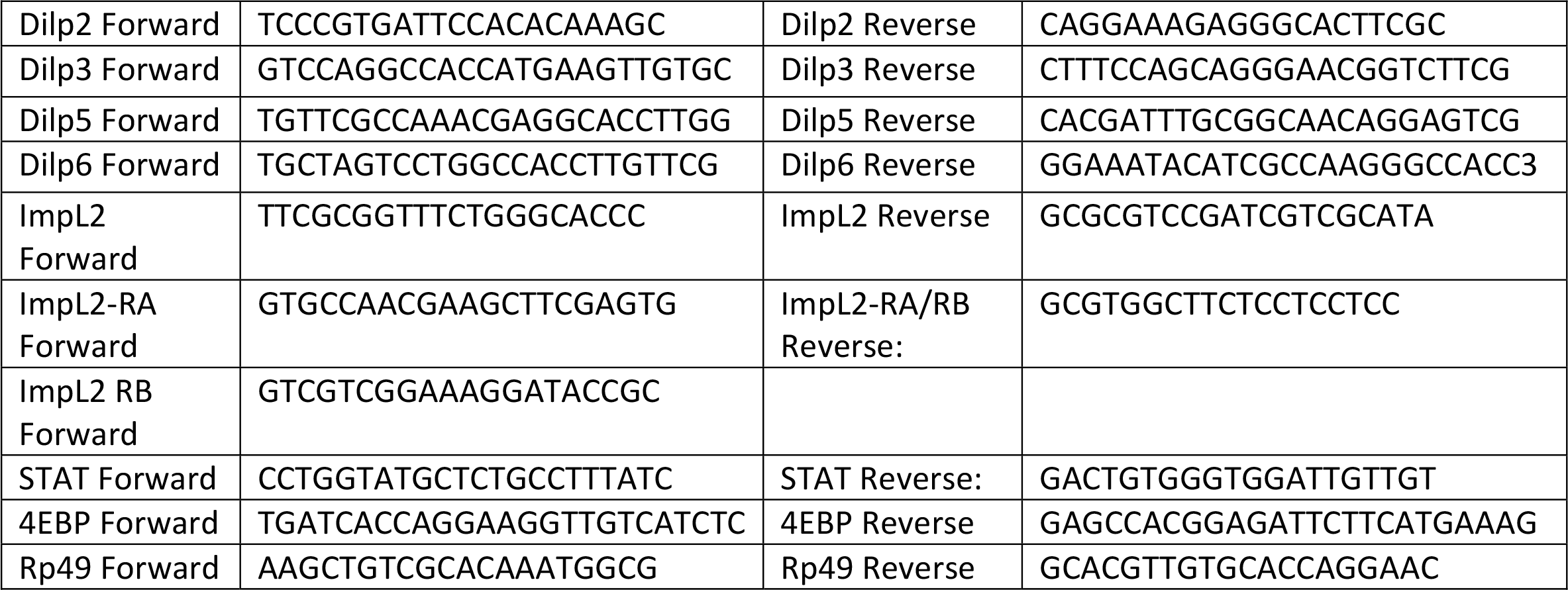

### Wasp infection

In all situations, cages containing approximately 200 virgin females and 100 males were allowed to lay eggs for 4 hours at 25 °C. Food plates containing embryos were incubated at 18 °C for 72 hours before being transferred to 25 °C for 18 hours. This was to restrict the induction of the Gal4 until late second instar larval phase, therefore circumventing any adverse effects during development. Early third instar larvae were transferred to a fresh food plate and *Leptopilina boulardi* were introduced. For high carbohydrate diet experiments, larvae were transferred to a high carbohydrate food plate at the point of infection. Wasps were allowed to infect for 40 to 45 minutes for standard infections and 15 to 20 minutes for weak infections. Wasps were then removed and infected larvae were incubated at 25 °C for a further 18 hours.

### Hemocyte count

Hemocytes counts were performed either 22 hours or 26 hours post infection. Individual third instar larvae were placed in 15 μl of PBS (Phosphate-buffered saline) and the cuticle was pierced using forceps to allow hemolymph to efflux. 10 μl of the hemolymph PBS mix was transferred to a hemocytometer (Neubauer). Samples were imaged using Leica ICC50 W (Leica, ICC50 W) Hemocytes that lay within the gridded area were counted, plasmatocytes and lamellocytes were differentiated by morphology. The total number of hemocytes per larvae was calculated by multiplying the count by 150. Difference between the genotypes was assessed using a one-way ANOVA.

### Resistance

Resistance to wasp infection was discerned by determining the viability of the Drosophila larvae or *Leptopilina boulardi* larvae. After weak infections, Drosophila larvae were left to pupate at 25 °C. Pupae were then dissected using forceps in PBS. The number of wasp larvae and melanized wasp eggs were counted in each larvae. Melanized eggs indicated an effective immune response and therefore likely survival of the Drosophila larvae. Live wasp larvae suggest the Drosophila larvae were unable to overcome infection and were out competed by *Leptopilina boulardi*. Statistical difference between genotypes was determined using a one-way ANOVA.

### Analysis of Metabolite levels: (Glucose GO kit)

30 third instar larvae were collected on ice, the cuticle was punctured using forceps and the hemolymph was transferred to a 1.5 ml Eppendorf tube containing 148 μL of 1x PBS. Tubes were centrifuged at 460 g, 4°C for 5 minutes. Supernatant was transferred to fresh Eppendorf tube and hemolymph was heat-inactivated at 70 °C for 15 min to degrade proteins. The hemolymph was centrifuged at 15 000 g, 4°C for 7 minutes and the supernatant was transferred and used for carbohydrate measurements. Glucose levels were obtained using a Glucose (GO) assay kit (Sigma-Aldrich) per manufacturer’s instructions.

Trehalose measurements were determined by adding trehalase (Sigma-Aldrich) to samples and incubating for 20 hours at 37 °C prior to measuring absorbance. Glycogen measurements were acquired from muscles of seven third instar larvae. The muscles attached to the cuticle of the animal were removed using forceps and added to a 1.5 ml Eppendorf tube containing 100 μL of 1x PBS. The tissue was homogenized and the centrifugation and heating steps were performed as for the hemolymph. Amyloglucosidase (Sigma-Aldrich) was added to each sample prior to incubation. Difference between the control and null mutants was assessed using a one-way ANOVA.

### FIM imaging

Larval locomotion was analyzed using frustrated total internal reflection-based (FIM) imaging (Risse *et al*., 2014). 10 third instar larvae were placed on an agar plate using a PBS soaked brush, following this larvae were recorded for 2 minutes 30 seconds at 10 frames per second. Tracking data was obtained using FIMTrack_v3.1.32.3 (http://fim.uni-muenster.de64) and analyzed using FIManalytics v0.10.1.2 and Excel. Statistical analysis was performed using FIManalytics and differences between genotypes were determined by a Mann-Whitney-U test. All experiments were conducted at room temperature. Accumulated distance is defined as the total length of larval trajectories per minute.

### Relative gene expression by qPCR

The muscle tissue (attached to the cuticle) of seven third instar larvae were dissected in PBS then homogenized in TRIzol reagent (Ambion). RNA isolation was performed using a Direct-zol RNA MicroPrep assay kit per manufactures instructions (ZYMO Research). Reverse transcription was carried out using PrimeScript RT Reagent Kit (TaKaRa) following manufactures instruction. mRNA expression for genes of interest were measured on FX 1000 Touch Real-Time Cycler (Bio-Rad) under the following conditions: 3 min denaturation at 95°C, 15 second amplification at 94°C, 15 seconds at 57°C, 15 seconds at 72°C for 40 cycles; melting curve analysis performed at 57 – 95°C/step 0.5°C. Data was analyzed using double delta Ct analysis, expressions was normalized to Ribosomal protein 49 (Rp49) expression. Relative expression of the target genes is presented as a fold change, with the control artificially set to 1. Difference between the control samples and mutants/knockdowns was assessed using ordinary one-way ANOVA.

### Bulk RNAseq analysis

Bulk RNAseq analysis was performed as described previously (preprint: Kazek *et al*., 2023).

## Acknowledgements

The authors acknowledge funding from the Grant Agency of the Czech Republic to TD (Project 20-09103S; www.gacr.cz) and EMBO (ALTF 23-2022). We thank Dr. Bruno Lemaitre, Dr. Hugo Stocker, Michele Crozatier and Bloomington Drosophila Stock Center for fly and wasp stocks. We thank Christian Klämbt for help with the FIM experiments. We thank to Lucie Hrádková for laboratory management and Marcela Jungwirthová for project management, and all members of Dolezal laboratory for their help with work.

## Figure legends

**Supplementary Figure 1:**
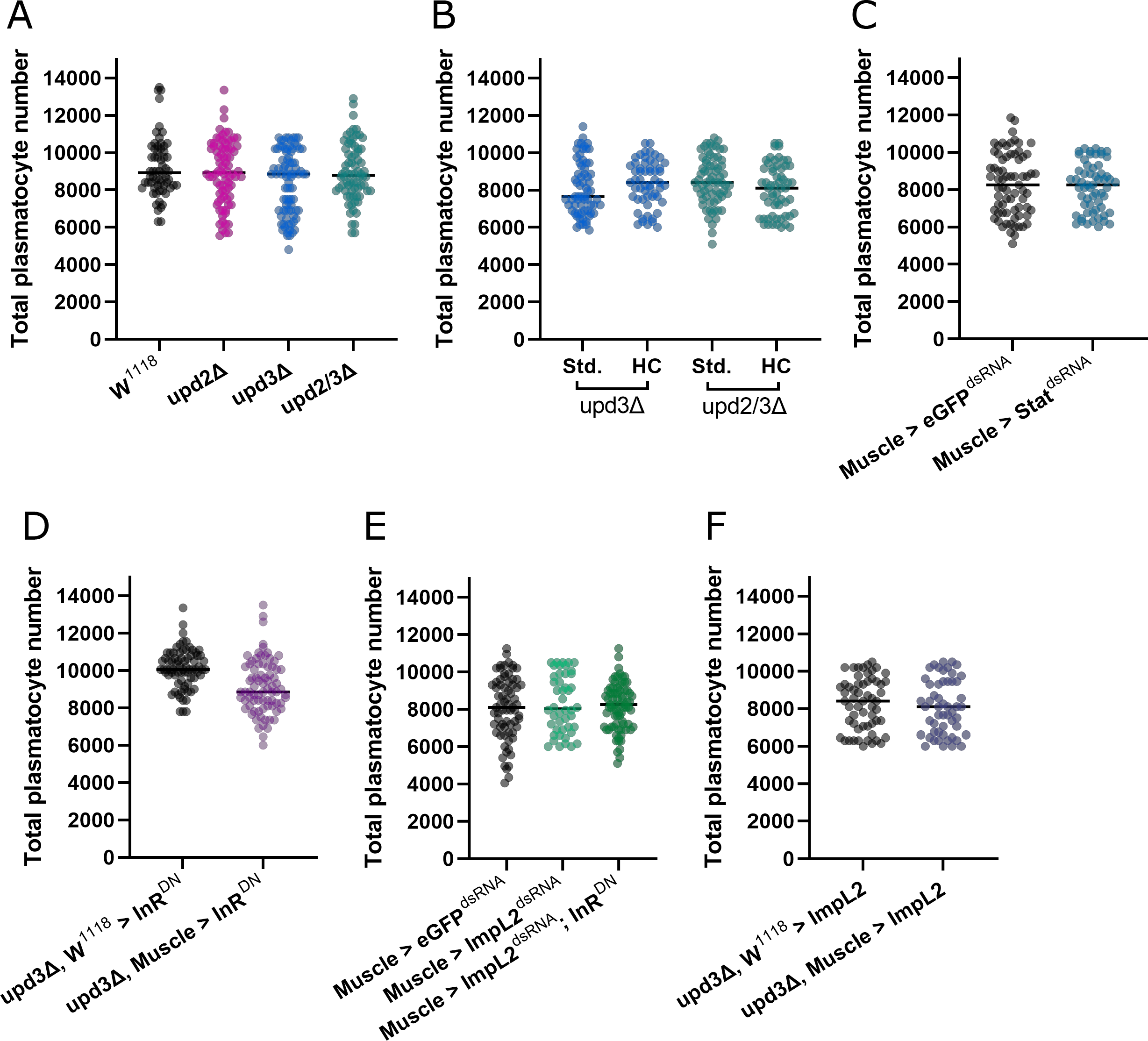
Plasmatocyte number is not effected by knockout or knockdown of Upd/JAK/STAT signaling Plasmatocyte count at 22 hpi for the following genotypes A: w^1118^ (control), upd2Δ, upd3Δ and upd2/3Δ (N=3, n=62-85). B: upd3Δ and upd2/3Δ on standard and high carbohydrate diet (N=3, n=56-67). C: Muscle > eGFP^dsRNA^ and Muscle > STAT^dsRNA^ (N=3, n=108-120). D: Upd3Δ, W^1118^ > InR^DN^ and Upd3Δ, Muscle > InR^DN^ (N=3, n=76-79). E: Muscle > eGFP^dsRNA^, Muscle > ImpL2^dsRNA^ and Muscle > ImpL2^dsRNA^ InR^DN^ (N=3, n=48-71). F: Upd3Δ, W^1118^ > ImpL2 and Upd3Δ, Muscle > ImpL2 (N=3, n=53-58). Each dot represents number of lamellocytes from an individual larva. There is no significant difference in the number of plasmatocytes. N represents individual experiments, n represents biological replicates.

**Supplementary Figure 2:**
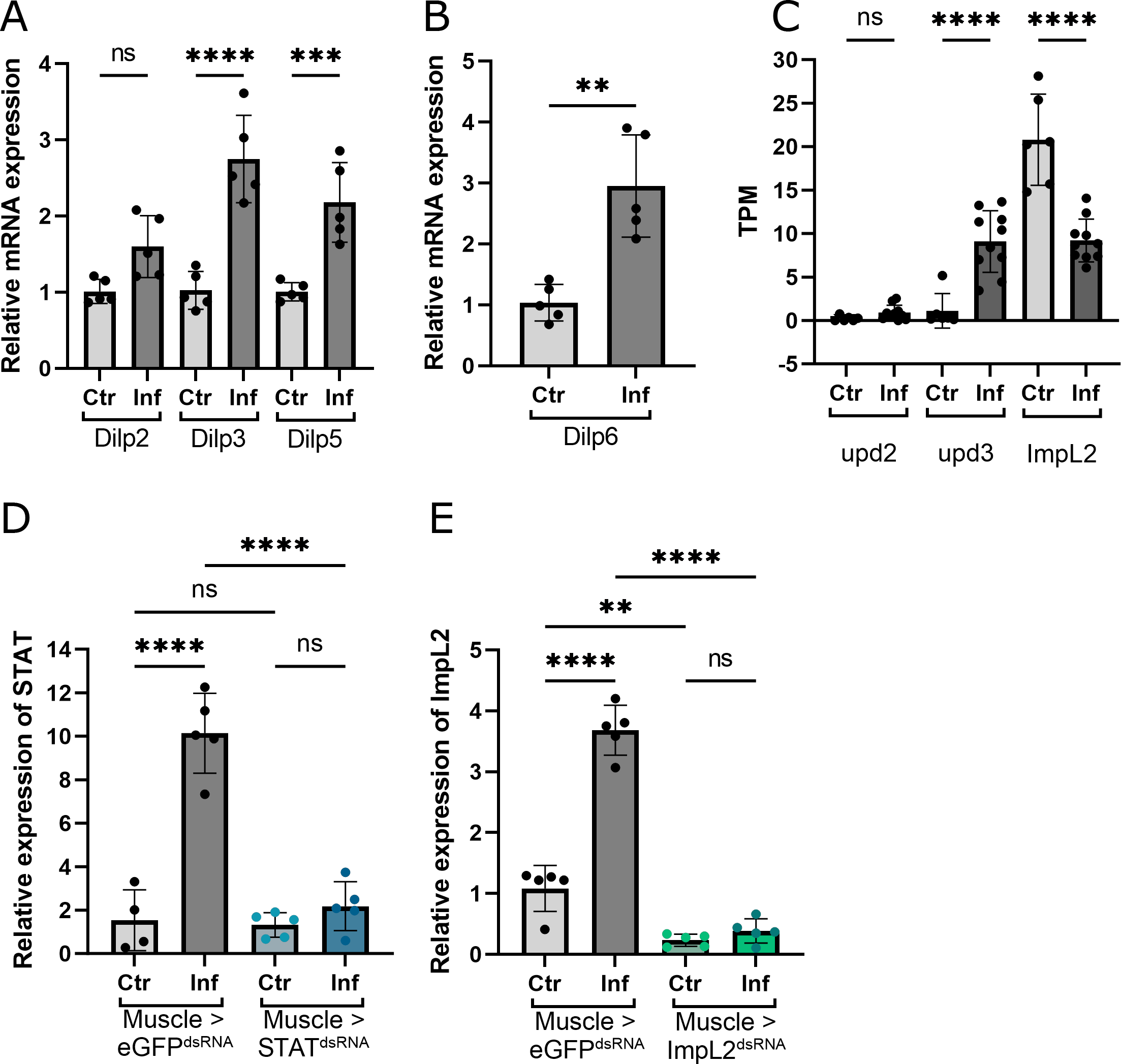
Expression of Dilps in the brain and fat body A: Relative expression of Dilp2, Dilp3 and Dilp5 in the central nervous system of control and infected third instar larvae 28 hpi (n=5). B: Comparative expression of Dilp6 in the fat body of control and infected animals (n=5). C: Expression of upd2, upd3 and ImpL2 (bulk RNAseq) in hemocytes with and without infection, Y axis shows transcripts per million (TPM) (n=6-10) D: Expression of STAT with and without infection in Muscle > eGFP^dsRNA^ and Muscle > STAT^dsRNA^ animals (n=4-5). E: Levels of ImpL2 in ImpL2 knockdown animals and controls with and without infection (n=5). Bars represent mean values, dots represent biological replicates. ns: no significant difference, ** p ≤ 0.01, *** p ≤ 0.001, **** ≤ 0.0001. N represents individual experiments, n represents biological replicates.

## References

Agaisse, H. et al. (2003) ‘Signaling Role of Hemocytes in Drosophila JAK/STAT-Dependent Response to Septic Injury’, Developmental Cell, 5(3), pp. 441–450. Available at: 10.1016/S1534-5807(03)00244-2.

Amoyel, M. et al. (2016) ‘Somatic stem cell differentiation is regulated by PI3K/Tor signaling in response to local cues’, Development, 143(21), pp. 3914–3925. Available at: 10.1242/dev.139782.

Bajgar, A. et al. (2015) ‘Extracellular Adenosine Mediates a Systemic Metabolic Switch during Immune Response’, PLOS Biology, 13(4), p. e1002135. Available at: 10.1371/journal.pbio.1002135.

Bajgar, A., Krejčová, G. and Doležal, T. (2021) ‘Polarization of Macrophages in Insects: Opening Gates for Immuno-Metabolic Research’, Frontiers in Cell and Developmental Biology, 9, p. 629238. Available at: 10.3389/fcell.2021.629238.

Ding, G. et al. (2021) ‘Coordination of tumor growth and host wasting by tumor-derived Upd3’, Cell Reports, 36(7), p. 109553. Available at: 10.1016/j.celrep.2021.109553.

Gurzov, E.N. et al. (2016) ‘The JAK/STAT pathway in obesity and diabetes’, The FEBS Journal, 283(16), pp. 3002–3015. Available at: 10.1111/febs.13709.

Honegger, B. et al. (2008) ‘Imp-L2, a putative homolog of vertebrate IGF-binding protein 7, counteracts insulin signaling in Drosophila and is essential for starvation resistance’, Journal of Biology, 7(3), p. 10. Available at: 10.1186/jbiol72.

Jünger, M.A. et al. (2003) ‘The Drosophila Forkhead transcription factor FOXO mediates the reduction in cell number associated with reduced insulin signaling’, Journal of Biology, 2(3), p. 20. Available at: 10.1186/1475-4924-2-20.

Kazek, M. et al. (2023) ‘Metabolism of glucose and trehalose by cyclic pentose phosphate pathway is essential for effective immune response in Drosophila’. bioRxiv, p. 2023.08.17.553657. Available at: 10.1101/2023.08.17.553657. [PREPRINT]

Krejčová, G. et al. (2021) ‘Macrophage-derived insulin/IGF antagonist ImpL2 regulates systemic metabolism for mounting an effective acute immune response in Drosophila’. bioRxiv, p. 2020.09.24.311670. Available at: 10.1101/2020.09.24.311670. [PREPRINT]

Liu, Ying et al. (2023) ‘Tumor Cytokine-Induced Hepatic Gluconeogenesis Contributes to Cancer Cachexia: Insights from Full Body Single Nuclei Sequencing’, bioRxiv, p. 2023.05.15.540823. Available at: 10.1101/2023.05.15.540823. [PREPRINT]

Lourido, F. et al. (2021) ‘Domeless receptor loss in fat body tissue reverts insulin resistance induced by a high-sugar diet in Drosophila melanogaster’, Scientific Reports, 11(1), p. 3263. Available at: 10.1038/s41598-021-82944-4.

Morin-Poulard, I., Vincent, A. and Crozatier, M. (2013) ‘The Drosophila JAK-STAT pathway in blood cell formation and immunity’, JAK-STAT, 2(3). Available at: 10.4161/jkst.25700.

Nappi, A.J. and Christensen, B.M. (2005) ‘Melanogenesis and associated cytotoxic reactions: applications to insect innate immunity’, Insect Biochemistry and Molecular Biology, 35(5), pp. 443–459. Available at: 10.1016/j.ibmb.2005.01.014.

Risse, B. et al. (2014) ‘FIM Imaging and FIMtrack: Two New Tools Allowing High-throughput and Cost Effective Locomotion Analysis’, Journal of Visualized Experiments : JoVE, (94), p. 52207. Available at: 10.3791/52207.

Russo, J. et al. (1996) ‘Insect immunity: early events in the encapsulation process of parasitoid (Leptopilina boulardi) eggs in resistant and susceptible strains of Drosophila’, Parasitology, 112 (Pt 1), pp. 135–142. Available at: 10.1017/s0031182000065173.

Shin, M. et al. (2020) ‘Subpopulation of Macrophage-Like Plasmatocytes Attenuates Systemic Growth via JAK/STAT in the Drosophila Fat Body’, Frontiers in Immunology, 11. Available at: https://www.frontiersin.org/articles/10.3389/fimmu.2020.00063.

Slaidina, M. et al. (2009) ‘A Drosophila Insulin-like Peptide Promotes Growth during Nonfeeding States’, Developmental Cell, 17(6), pp. 874–884. Available at: 10.1016/j.devcel.2009.10.009.

Smith, R.L. et al. (2018) ‘Metabolic Flexibility as an Adaptation to Energy Resources and Requirements in Health and Disease’, Endocrine Reviews, 39(4), pp. 489–517. Available at: 10.1210/er.2017-00211.

Son, H.G. et al. (2019) ‘Age-dependent changes and biomarkers of aging in Caenorhabditis elegans’, Aging Cell, 18(2), p. e12853. Available at: 10.1111/acel.12853.

Straub, R.H. (2014) ‘Insulin resistance, selfish brain, and selfish immune system: an evolutionarily positively selected program used in chronic inflammatory diseases’, Arthritis Research & Therapy, 16(2), p. S4. Available at: 10.1186/ar4688.

Teleman, A.A., Chen, Y.-W. and Cohen, S.M. (2005) ‘4E-BP functions as a metabolic brake used under stress conditions but not during normal growth’, Genes & Development, 19(16), pp. 1844–1848. Available at: 10.1101/gad.341505.

Terry, N.A. et al. (2006) ‘Novel regulators revealed by profiling Drosophila testis stem cells within their niche’, Developmental Biology, 294(1), pp. 246–257. Available at: 10.1016/j.ydbio.2006.02.048.

Tokusumi, T. et al. (2009) ‘Characterization of a Lamellocyte Transcriptional Enhancer Located within the misshapen Gene of Drosophila melanogaster’, PLoS ONE, 4(7), p. e6429. Available at: 10.1371/journal.pone.0006429.

Wang, A., Luan, H.H. and Medzhitov, R. (2019) ‘An evolutionary perspective on immunometabolism’, Science, 363(6423), p. eaar3932. Available at: 10.1126/science.aar3932.

Wang, L., Karpac, J. and Jasper, H. (2014) ‘Promoting longevity by maintaining metabolic and proliferative homeostasis’, The Journal of Experimental Biology, 217(1), pp. 109–118. Available at: 10.1242/jeb.089920.

Yang, H. et al. (2015) ‘JAK/STAT signaling in Drosophila muscles controls the cellular immune response against parasitoid infection’, EMBO reports, 16(12), pp. 1664–1672. Available at: 10.15252/embr.201540277.

Yang, H. and Hultmark, D. (2017) ‘Drosophila muscles regulate the immune response against wasp infection via carbohydrate metabolism’, Scientific Reports, 7, p. 15713. Available at: 10.1038/s41598-017-15940-2.

Yu, S. et al. (2022) ‘Drosophila Innate Immunity Involves Multiple Signaling Pathways and Coordinated Communication Between Different Tissues’, Frontiers in Immunology, 13, p. 905370. Available at: 10.3389/fimmu.2022.905370.

